# Assessing tensor decomposition quality of immune profiling data from a dictionary learning perspective

**DOI:** 10.64898/2026.07.03.736447

**Authors:** Anna Konstorum, Jian Xing, Shuchin Aeron, Misha Kilmer, Steven H. Kleinstein

**Affiliations:** Center for Computing Sciences, Institute for Defense Analyses, U.S.A.; Department of Pathology, Yale School of Medicine, New Haven, Connecticut, U.S.A.; Department of Electrical and Computer Engineering, Tufts University, U.S.A.; Department of Mathematics, Tufts University, U.S.A.; Department of Immunobiology, Yale School of Medicine, New Haven, Connecticut, U.S.A.; Department of Biomedical Informatics and Data Science, Yale School of Medicine, New Haven, Connecticut, U.S.A.; Program in Computational Biology and Biomedical Informatics, Yale University, New Haven, Connecticut, U.S.A.

## Abstract

Systems-level immune profiling data arising from longitudinal studies of vaccination or infection has an inherent multi-index array structure. While tensor decomposition of such datasets has gained popularity, choosing a rank and trial for a decomposition is not straightforward. We show that taking into account the experimental data model can inspire the development of new metrics to assess the quality of a Non-negative CANDECOMP/PARAFAC (NCPD) decomposition, and can thus be used to choose a rank and trial for the decomposition. Moreover, we show how framing the results via a dictionary learning framework can better enable interpretation of the components of the decomposition.

## 1 Introduction

Systems-level immune profiling datasets increasingly consist of complex study designs that can involve, for example, omics data collected from multiple tissues longitudinally [1–3]. Although these studies apply a range of techniques to analyze high-dimensional immunological data, analyses are typically performed on two-dimensional representations (e.g., gene-by-sample), which require separating the data by time-point, tissue, or modality and lead to largely time-point- or modality-specific insights. Consequently, higher-order structure, such as coordinated patterns spanning individuals, genes, time, and tissues, is not explicitly modeled, necessitating the development of novel mathematical approaches that account for the inherently multi-dimensional nature of these datasets.

Multi-index (tensor) frameworks can serve as a valuable tool to organize and analyze datasets with complex study designs [4]. For example, a study examining gene expression over time in a set of participants exposed to a treatment could naturally be structured as a participant × gene × time tensor; flattening the data into a matrix would result in a loss of information (e.g. reformatting the data into a gene *×* (participant × time) matrix results in a loss of temporal information).

Tensor decomposition can deconstruct a complex multi-dimensional dataset into a small number of simple, interpretable patterns across each dimension. Thus, a decomposition of a tensor that includes experimental axes such as participants, genes (or other features), and time can result in efficient characterization of multi-index patterns of variation in the datasets, analogous to Principal Components Analysis (PCA) or Non-negative Matrix Factorization (NMF) for two-dimensional datasets. In the life sciences, some applications of tensor decompositions have included the identification of multi-scale patterns in neural circuits during learning [5], tissue-specific gene expression phenotyping linked to genomic variation [6], *Mycobactaerium tuberculosis* subtyping [7], and systems-serology subtyping [8]. Nevertheless, several challenges remain in applying tensor decompositions robustly to biological time-course datasets, such as choosing the number of patterns in the decomposition (rank), and the best decomposition for a given rank (trial), as well as assessing model quality in light of interpretability [9].

The non-negative CANDECOMP/PARAFAC tensor decomposition (NCPD) extends the idea of non-negative matrix factorization (NMF) to multi-dimensional data by representing a tensor as a sum of non-negative rank-one components. For a participant *×* feature *×* time tensor, each component captures a coordinated pattern involving a subset of participants, a subset of features, and an associated temporal trajectory. As such, NCPD provides a natural framework for identifying interpretable patterns in longitudinal immune profiling data. While uniqueness and existence results have been established for NCPD [10], it has also been observed that a decomposition with minimal reconstruction error may not provide the best match to an underlying data model [9]. Therefore, in order to apply NCPD effectively to immune profiling data, it is important to develop measures of decomposition quality that account not only for reconstruction accuracy, but also for model match and component-level interpretability.

In this work, we introduce an extension of NMF for a third order tensor with a temporal mode, the *Feature Canonical Trajectory* (FCT) tensor dictionary learning framework, in which the data for each participant are modeled as a weighted combination of low-rank matrix-valued elements that capture feature behavior over time. This formulation provides a generative interpretation of the NCPD in context of immune profiling data, in which the components correspond directly to interpretable building blocks that represent both feature and temporal structure. This framework aids in the interpretation and analysis of the decomposition by providing a higher-dimensional analogue to the concept of gene or feature modules that contribute to the observed response. Furthermore, we use this framework to develop data model-appropriate component-level measures to help choose a rank and trial for an NCPD of immune profiling data. We show, using real and synthetic datasets, how the new component-level measures can be used in concert with established metrics to evaluate and choose a decomposition that improves model match and interpretability (Figure 1).

**Fig 1.**
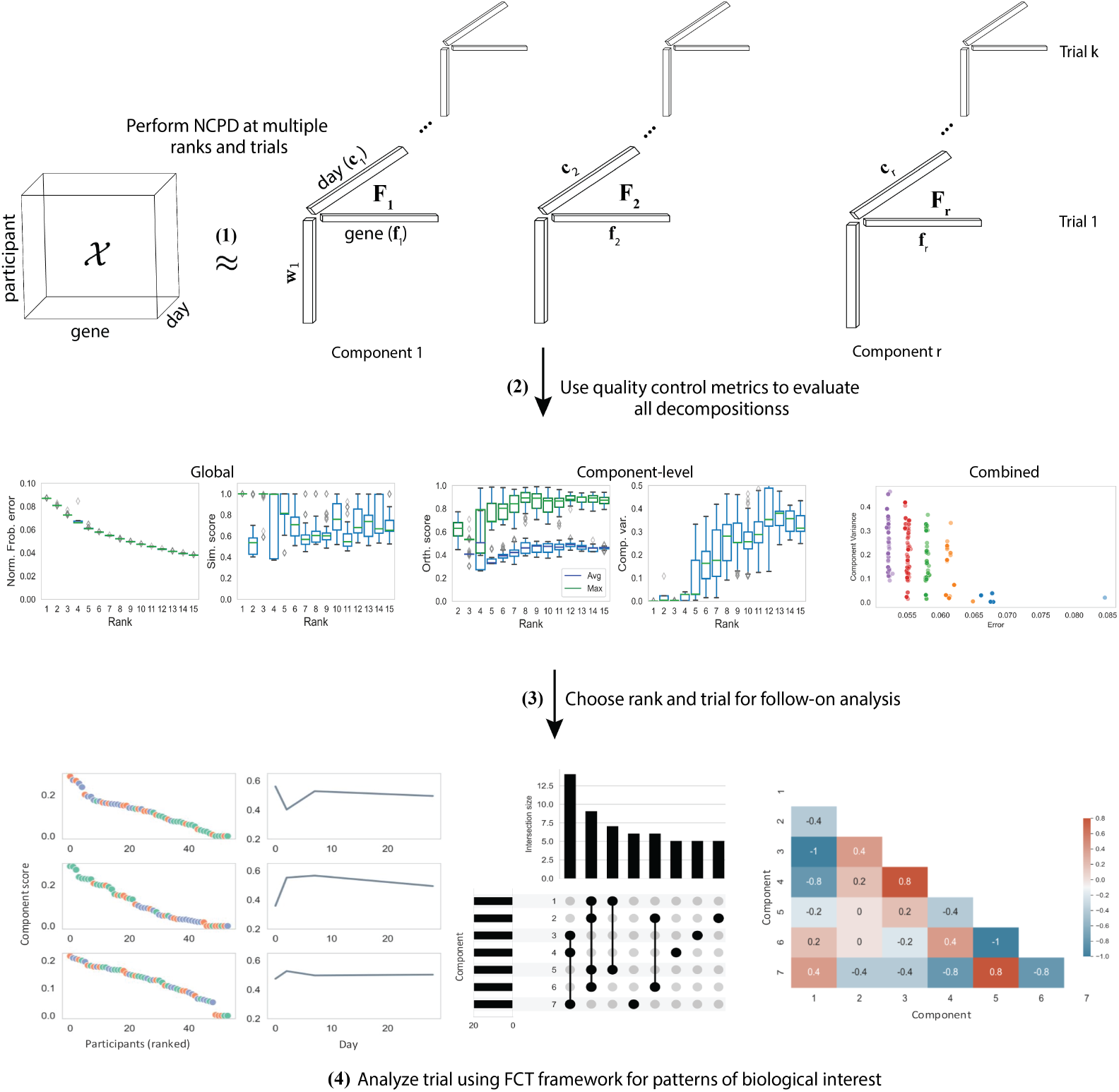
Overview of pipeline to choose and evaluate a CP decomposition of immune profiling data. Steps include (1) perform decompositions over a range of ranks, and multiple trials per rank, (2) use global and novel component-level quality control metrics to evaluate decomposition performance, (3) choose a rank and trial for follow-on analysis, and (4) use the FCT tensor dictionary learning framework to evaluate and interpret results.

## 2 Results

Many modern immunological studies generate high-dimensional, longitudinal datasets by measuring multiple molecular features, such as genes, proteins, or metabolites, across a cohort of participants over a series of time points. We refer to the collection of measurements for all features across all time points for a given sample as its *feature profile*. While our framework is general and could be applied to transcriptomics, proteomics, or metabolomics data, in this work we focus on transcriptomic profiling experiments, where gene expression is measured over time. Such data can be naturally organized as a third-order participant × feature × time tensor, 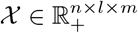, where *n* participants are measured for *l* features over *m* time points (Figure 1).

Conceptually, each participant’s feature profiles can be thought of as a combination of basic building blocks, or dictionary elements, where each element captures a coordinated set of features that follow a shared temporal pattern. We represent each horizontal slice of *X* — corresponding to the feature profiles of a single participant over time, as a sum of these rank-one matrix dictionary elements, which we term *Feature Canonical Trajectories* (FCTs). Each FCT is further decomposed into the outer product of a trajectory vector (*Canonical Trajectory*, CT), describing temporal dynamics, and a feature vector (*Canonical Feature-set*, CF), describing the subset of features that exhibits this trajectory. This construction naturally leads to a generative model for a nonnegative CP decomposition (NCPD) of the data tensor *X*.

### 2.1 Notation and Preliminaries

Scalars, vectors, matrices, and tensors are represented by lower-case letters (*a*), lower-case bold-face letters (**a**), uppercase bold-face letters (**A**), and calligraphic uppercase letters (*A*), respectively. The number of dimensions of a tensor is referred to as the *order*. Column *j* of a matrix **A** is written compactly as **a**_*j*_, the (*i, j*) entry of **A** is written as *a*_*ij*_, and the *i*th entry of a vector **a** is written as *a*_*i*_. A *n*th element of a sequence of matrices (vectors) is denoted by **A**^(*n*)^ (**a**^(*n*)^).

A *slice* is a two-dimensional tensor subarray that results from fixing all but two indices of a tensor. For a third order tensor *X*, a slice can be denoted as *Xi*_::_ (horizontal slice), *X*_:*j*:_ (lateral slice), or *X*_::*k*_ (frontal slice), where the colon notation indicates that the index in that position takes on all values. For example, if *X* ∈ ℝ^*n×l×m*^ is a (participant) by (gene) by (time) tensor, then *Xk*_::_ ∈ ℝ^1*×l×m*^ corresponds (after reshaping) to an *l × m* (gene) by (time) matrix representing the *k*th horizontal slice. In this study, we are particularly interested in the horizontal slices, which each represent a participant. Therefore, for participant *k*, we reshape the tensor *X*_*k*_:: into the matrix **X**_*k*_ as follows,

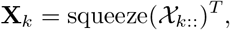

where the *squeeze* function removes the dimensions of length one.

A *fiber* is a one-dimensional tensor subarray (vector) such that all indices except for one are fixed. For a third order tensor *X*, the fiber associated with fixing the first two indices is referred to as a *tube fiber*, and is denoted *X*_*ij*:_.

We introduce the operation `vec` () to move between a tube fiber and a vector **t** ∈ ℝ^*m*^ as follows,

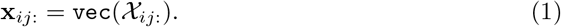

A *component* is an outer product of two or more vectors (also termed a rank-one matrix or tensor). Since we will be analyzing the components of a decomposition of a third-order tensor, we formally define a rank-one third-order tensor.

#### Definition 1

*A third order tensor X* ∈ ℝ^*n×l×m*^ *is* rank one *if it can be written as an outer product of three vectors, X* = **a** ◦ **b** ◦ **c**, *where* **a** ∈ ℝ^*n*^, **b** ∈ ℝ^*l*^, **c** ∈ ℝ^*m*^ *and* ◦ *denotes the vector outer product*.

By definition, the (*i, j, k*) entry of a rank-one third order tensor *X* is *X*_*ijk*_ = *a*_*i*_*b*_*j*_*c*_*k*_.

For a more extensive discussion of notational terminology associated with tensors, see [4].

#### 2.1.1 Non-negative CP decomposition (NCPD)

For a non-negative matrix, 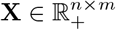, non-negative matrix factorization (NMF) finds a rank *r* decomposition,

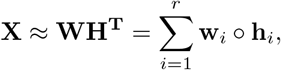

where 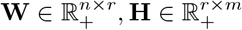, and ◦ denotes the vector outer product.

In non-negative CANDECOMP/PARAFAC Decomposition (NCPD) the concept in NMF of decomposing a matrix into a non-negative sum of rank-one components is extended to tensors. For a third order tensor 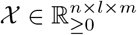, the associated NCPD decomposition of rank *R* is as follows,

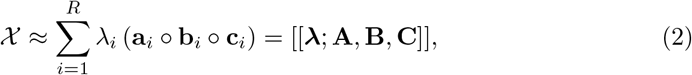

where||**a**_*i*_||_2_=||**b**_*i*_||_2_ = ||**c**_*i*_||_2_ = 1 for all *i*. The matrices, 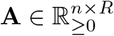, 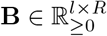, 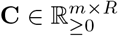 have **a**_*i*_, **b**_*i*_, **c**_*i*_ as their columns, respectively, and are termed *factor matrices*, and 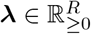 is a vector of normalization constants. The components are ordered in decreasing magnitude of *λ*_*i*_. The compact representation of the NCPD on the right-hand side of Equation 2 is termed the *Kruskal notation*, and can be extended in a straightforward manner to higher orders [4].

#### 2.1.1 The Feature Canonical Trajectory framework for time-course data

Given a participant × feature × time tensor 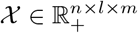, we begin by defining the building blocks of an FCT, Canonical Trajectories and Canonical Feature-sets. To ground the discussion, we use a synthetic dataset (described fully in the Methods) generated using the FCT framework. This dataset is a nonnegative participant *×* feature *×* time tensor representing a cohort of three groups of individuals (healthy young, healthy older, and immunocompromised) who are vaccinated on day 1, with blood transcriptomic data collected at baseline (day 1) and at three subsequent time points. Each participant exhibits a cohort-specific response to vaccination, characterized by distinct subsets of features (referred to as *feature-sets*) that follow different temporal patterns. These response patterns enable characterization of participants based on their individual response profiles (Figure 2).

**Fig 2.**
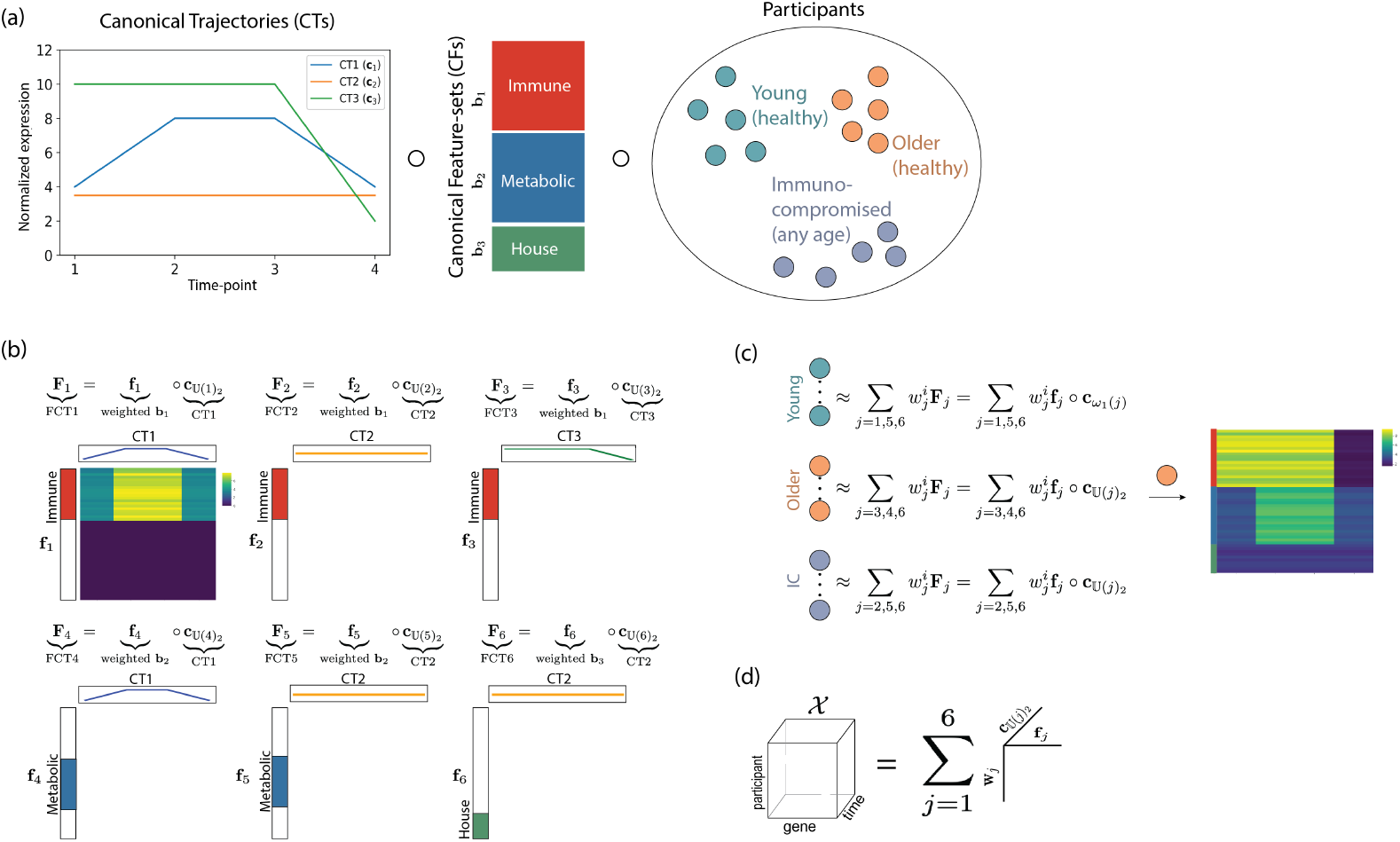
Synthetic data generated from the FCT model. (a) The data is composed of three Canonical Trajectories (CTs) and three Canonical Feature sets (CFs) that can follow any of these CTs; participants come from three groups: healthy Young, healthy Older, and Immunocompromised of any age; (b) Feature Canonical Trajectories (FCTs) are formed from the outer product of a CT and CF; (c) each group of participants correspond to the weighted sum of different FCTs; (d) the tensor is composed of the sum of components 1, 2, .., 6 that correspond to the outer product of the participant weights (**w**_*j*_ ) with the weighted CF (**f**_*j*_ ) and 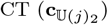 that correspond to FCT *j* (**F**_*j*_ ).

##### Definition 2

*A trajectory* 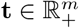 *of X is a vector that represents each tube fiber X*_*ij*:_ *as* **x**_*ij*:_ = *vec*(*X*_*ij*:_).

Note that there is a trajectory **x**_*ij*:_ associated with each participant *i* and feature *j*.

##### Definition 3

*A Canonical Trajectory (CT)*, 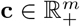, *is a temporal pattern that is similar to a set of trajectories, i*.*e*.

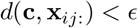

*for some ϵ, distance measure d*(*·, ·*), *i* ∈ *I* ⊆ [1, 2, …, *n*], *and j* ∈ *J* ⊆ [1, 2, …, *l*].

##### Definition 4

*A Canonical feature-set (CF)*, **g** ∈ *{*0, 1*}* ^ℓ^, *for a given set of indices J* ⊆ [1, 2, …, ℓ], *is defined as follows*,

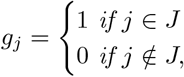

*for j* ∈ *J* ⊆ [1, 2, …, ℓ].

A canonical feature-set represents as subset of features that show a distinct response over time, as will be shown.

We use these building blocks to formulate a model that extends the concept that a participant *×* feature *×* time dataset can be a ‘weighted sum of its parts’ analogous to NMF. We model a participant *i*, represented as the matricized horizontal slice of *X* (**X**^*i*^) as a weighted combination of CFs that correspond to specific CTs. We consider the following unit that is analogous to a topic or hidden variable in NMF.

##### Definition 5

*A Feature Canonical Trajectory (FCT)* 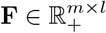, *is an outer product of a weighted CF with a CT*.

A Feature Canonical Trajectory (FCT) represents a set of features (CF) that have trajectories similar to one canonical trajectory (CT) over a time-course. The CTs for the synthetic data comprise three trajectories (Figure 2(a)): a trajectory that is constant and then drops at the last time-point (**c**_1_), a constant trajectory (**c**_2_), and a rise-and-fall trajectory (**c**_3_). The CFs correspond to three feature-sets: immune, metabolic, and house-keeping genes (Figure 2(a)). FCTs are formed by taking outer products between different CTs and CFs. For example, FCT1 (**F**_1_) is formed as the outer product of the immune-gene CF with the rise and fall CT ((Figure 2(b)). This rank-one matrix thereby represents immune genes as having a rise-and-fall trajectory. FCT2 (**F**_2_) on the other hand, the outer product of the immune gene CF and the constant CT, generates a matrix whereby immune genes are all constant.

The slice **X**_*i*_ corresponding to each participant is represented as a combination of FCTs. Older participants in the synthetic data are represented as a sum of FCTs 3,4, and 6 (Figure 2(c)), which correspond to immune genes that drop towards the end of the time-course (**F**_3_), metabolic genes with a rise-and-fall trajectory (**F**_4_) and constant levels of house-keeping genes (**F**_6_). The FCT framework is thus analogous to the topic or hidden variable as presented in NMF by considering each participant as a weighted combination of FCTs (Figure 2(d)).

### 2.3 Component-level quality control measures for rank and trial choice

We recall that the rank of a decomposition is the number of components it contains. Since an NCPD is, in general, not deterministic, it can yield different results even when computed at the same rank. We refer to each resulting decomposition at a given rank as a *trial*. Our goal is to develop measures for selecting both the rank and trial of an NCPD for immune profiling data that maximize the *interpretability* of the decomposition. We consider a decomposition to be interpretable when each trajectory is assigned to a single component and follows the temporal pattern represented by that component.

Component-level quality control measures are motivated by the data model for the NCPD of immune profiling data as articulated by the FCT framework: each component consists of an outer product of a canonical trajectory (CT), a canonical feature-set (CF), and a vector corresponding to participants in whom the features defined by the CF follow the associated CT. Therefore, one can assess the quality of a decomposition at the component level by evaluating whether the participants identified by a component express the features in the CF according to the pattern represented by the CT. Here we introduce two new measures, *{*1, 2*}-orthogonality* and *component variance*, that quantify this notion.

The *{*1, 2*}*-orthogonality metric evaluates the extent to which components are distinct by measuring overlap in participant–feature assignments, low values indicate that each trajectory is associated with a single component, improving interpretability. We refer to decompositions with low *{*1, 2*}*-orthogonality as having good *component separation*. In contrast, the component variance metric assesses how well each component’s CT represents the data assigned to it by quantifying the similarity between high-weight trajectories and the corresponding CT. Low component variance indicates that the component provides a faithful representation of the underlying data, we refer to such decompositions as having good *internal consistency*. Together, these metrics capture complementary aspects of model quality, namely separation between components and internal consistency within components.

The *{*1, 2*}*-orthogonality and component variance scores can be used in tandem with global measures, to assess component-level quality of decompositions. Global measures evaluate the overall quality of a decomposition, either in terms of its reconstruction accuracy with respect to the original data or its consistency with other decompositions.

While the ideal decomposition will have low scores for both component-level measures, low scores in either can still provide meaningful insight into model quality, as these measures capture aspects such as component separation and internal consistency that may be missed by global measures.

### 2.4 Component-level metrics showed a fine-grained quality resolution of NCPD on synthetic data

We first investigate whether component-level QC measures provide information beyond conventional global measures using a synthetic dataset with known ground truth (Figure 2). We evaluate decompositions across different ranks (number of components) and trials (independent runs at a fixed rank) to understand how these measures provide complementary information.

Using only global measures (Figure 3(a)), we observe that error approaches zero at ranks 6–7 and similarity declines beyond rank 7, suggesting that the true rank lies near *R* = 6. Nevertheless, these measures do not yield a conclusive result for selecting between candidate ranks or distinguishing between trials. Component-level measures provide additional insight: orthogonality increases with both under- and over-factoring, while component variance increases with over-factoring. Visualizing trials from ranks 5–7 in error × orthogonality × component variance space shows that rank-6 trials cluster at low values of all three measures, whereas rank-7 trials, despite low error, exhibit variability in orthogonality and component variance. This demonstrates that global measures alone may be insufficient for selecting a rank and trial, whereas component-level measures provide additional insight into the structure of the decomposition and can thus be used to aid in rank and trial selection.

**Fig 3.**
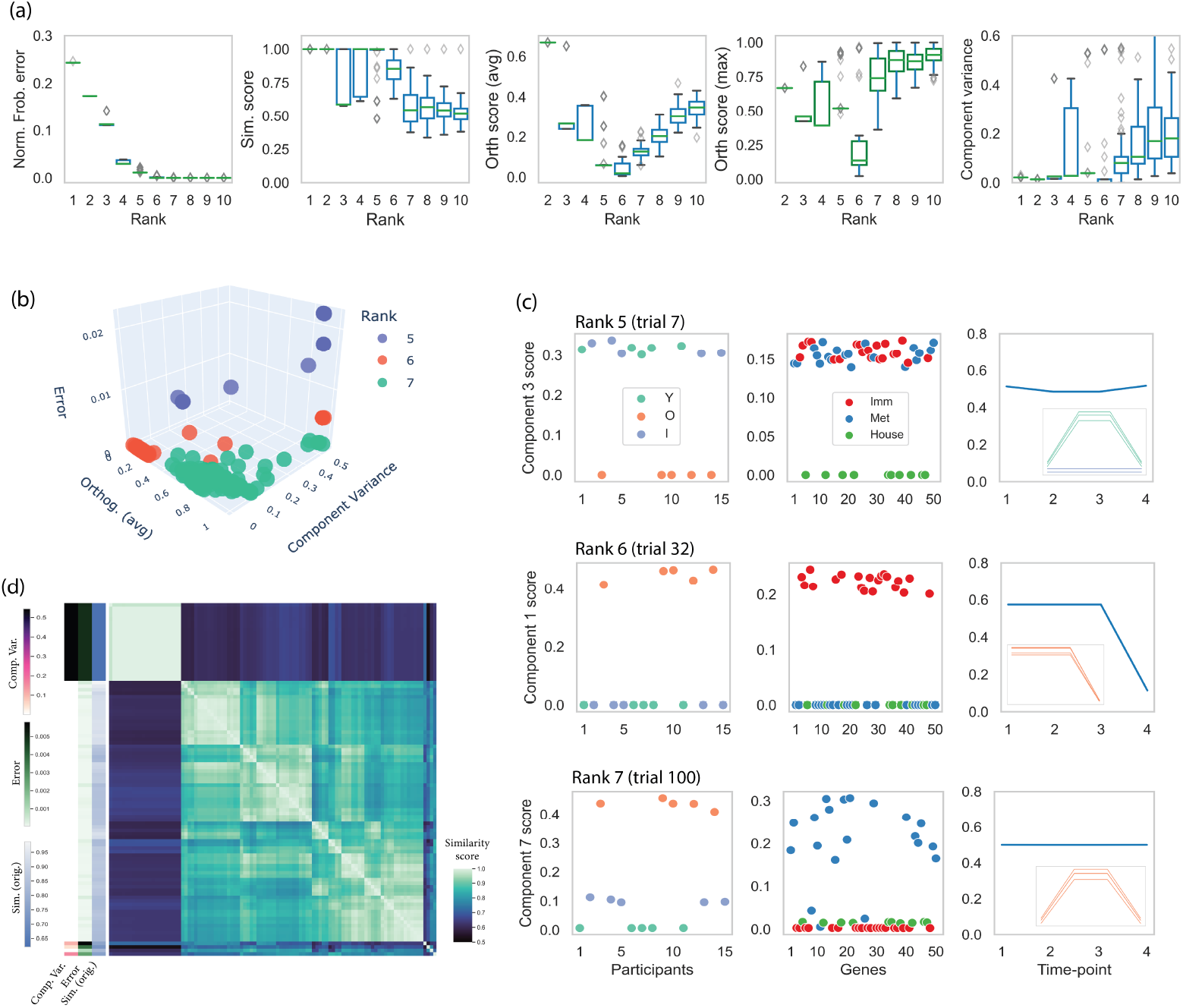
Component-level measures on synthetic data show utility in identifying interpretable NCPD trials. (a) Global and component-level quality results for NCPD of synthetic data with ranks *R* = 1 *−* 10, 100 trials per rank. The true rank is *R* = 6. (b) Plot of error by orthogonality (avg.) by component variance for trials of ranks 5-7. (c) Plotting trials from decompositions of ranks 5-7 with the lowest error. (d) Heatmap of similarity across rank 6 trials.

We consider a decomposition to be uninterpretable when its components are not well-separated or internally consistent, i.e. when trajectories are assigned to multiple components or do not follow the temporal pattern represented by their assigned component. To investigate how a trial that is uninterpretable may behave, we plotted components from the lowest error trials for each of ranks 5-7 and observed that the trials from ranks 5 and 7 contain components where the CTs do not correspond to the actual trajectories of the top scoring participants and genes (Figure 3(c)). These results show that error is not sufficient to determine the interpretability, and thus quality, of a decomposition, and component-level measures must be taken into account when choosing a rank and trial. Moreover, even for the case of rank 6 trials, we observe a not insignificant number of trials with higher error, component variance, and lower similarity to the original model (Figure 3(d)). Although these trials are all higher in error, and thus error alone could exclude them from subsequent analysis, these results show that even for the true rank decomposition, the higher error trials will result in lower interpretability of components, thus necessitating multiple trials for choice of downstream analysis.

A property of the FCT framework is that the resulting decompositions are not unique, meaning that the same underlying biological patterns can be represented by different combinations of components across trials. To illustrate this, we consider the rank-6 trial with lowest error (which also has low component variance and orthogonality scores), for which components 1–4 closely match the ground truth components (Supplementary Figure 1). However, the final two components differ in both participant and gene factor weights (Figure 4(a,b)). This can be explained by the fact that CTs and CFs can combine in multiple ways to form different FCTs that, taken together, encode the same underlying structure (Supplementary Figure 2).

**Fig 4.**
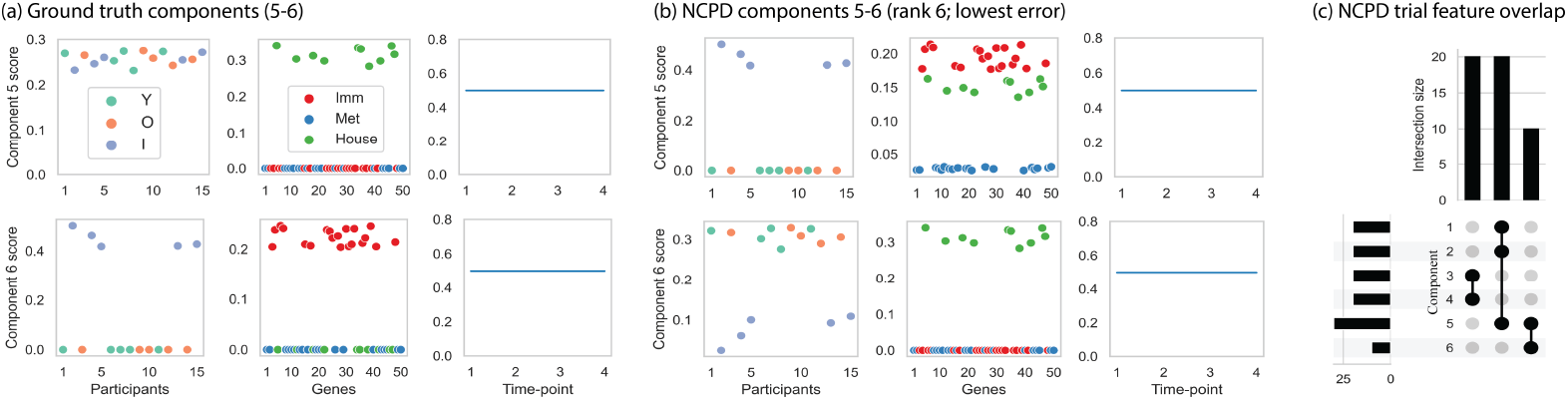
The difference between (a) ground-truth components 5 and 6 of the synthetic data model and (b) the corresponding components from the selected rank-6 trial is explained by a reorganization of the CFs across the components, resulting in an equivalent model. (c) UpSet plot of overlap among top-scoring features across components.

Importantly, this non-uniqueness does not diminish the interpretability of the decomposition. An intersection analysis of gene factor weights across components shows that component 5 shares one set of genes with components 1 and 2, and another set with component 6 (Figure 4(c)). These sets correspond to immune-related and housekeeping genes, respectively. Thus, even when individual components differ across trials, the CFs corresponding to the underlying data model can be recovered for decompositions with strong component-level QC scores (Supplementary Materials).

### 2.5 The FCT framework on an NCPD of real-world data can identify novel biological patterns

We next apply global and component-level QC measures to a real-world platelet transcriptomic dataset collected longitudinally from individuals before and after influenza vaccination, forming a participant *×* gene *×* time tensor. In contrast to the synthetic setting, the true rank is unknown, and our goal is to use QC measures to guide the selection of a rank and trial that yields an interpretable decomposition.

We first evaluated decompositions across ranks *R* = 1 − 15. Reconstruction error decreased smoothly with increasing rank, while similarity became more variable for ranks *R* > 3. The average and maximum orthogonality scores increased through approximately *R* = 8 and then began to level off; component variance followed a similar pattern, with its largest changes occurring before *R* = 8 (Figure 5(a)). Together, these trends suggested that ranks 4 - 8 contained the most relevant range for further evaluation.

**Fig 5.**
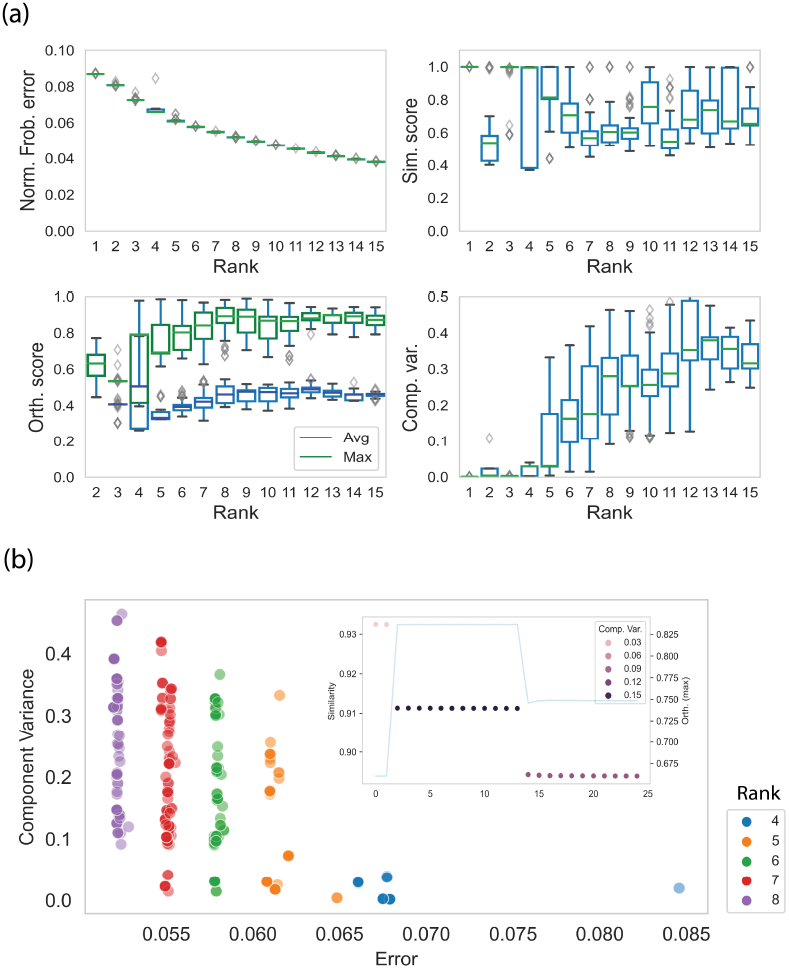
Quality analysis of NCPD decompositions for real-world data. (a) Global and component-level measures for decompositions of ranks 1–15, with 100 trials per rank. (b) Component variance versus reconstruction error for ranks 4–8. The inset shows the similarity scores (left axis, scatter) and *{*1, 2*}*-orthogonality scores (right axis, line) between the rank-5 trial with the lowest component variance and all rank-6 trials.

Within this range, plotting component variance against reconstruction error revealed a tradeoff: as rank increased, error tended to decrease, but component variance became more variable across trials, particularly at *R* = 8 (Figure 5(b)). We therefore used the component-level metrics to compare candidate trials across adjacent ranks. Starting from the rank-5 trial with the lowest component variance, we compared its similarity to all rank-6 trials and identified a rank-6 trial with high similarity, low component variance, and low orthogonality (Figure 5(b), inset). Applying the same logic to rank 7, we selected the rank-7 trial with the lowest component variance for downstream analysis because it remained highly similar to the preceding selected trials while adding another interpretable component (Supplementary Figure 3). These results demonstrate that QC-guided selection can identify candidate decompositions with improved interpretability beyond what is suggested by global measures alone.

To assess the biological relevance of the selected decomposition, we examine the components of the rank-7 trial and their associations with participant characteristics and known pathways. The majority of the components are correlated with age (Figure 6(a), Supplementary Table 1), and the top CFs identified using an intersection analysis are associated with pathways related to translation (components 3,4, and 7) and platelet activation (components 1,2,5,6). Interestingly, these two pathway programs were also identified in the preceding analysis [11], although that analysis identified only two components related to translation and three related to platelet activation. In contrast, the selected rank-7 decomposition identifies three components related to translation and four related to platelet activation. Thus, using the new quality control metrics, we were able to identify a higher-rank trial and additional components, such as component 6, which is more strongly associated with younger and older community-dwelling participants than with skilled nursing facility (SNF) participants and captures a platelet activation program that continues to increase throughout the time course. Moreover, the original analysis did not identify any components associated with vaccine responsiveness, whereas in the new analysis, components 1 and 6 both showed significant correlation with vaccine responsiveness (Supplementary Table 1).

**Fig 6.**
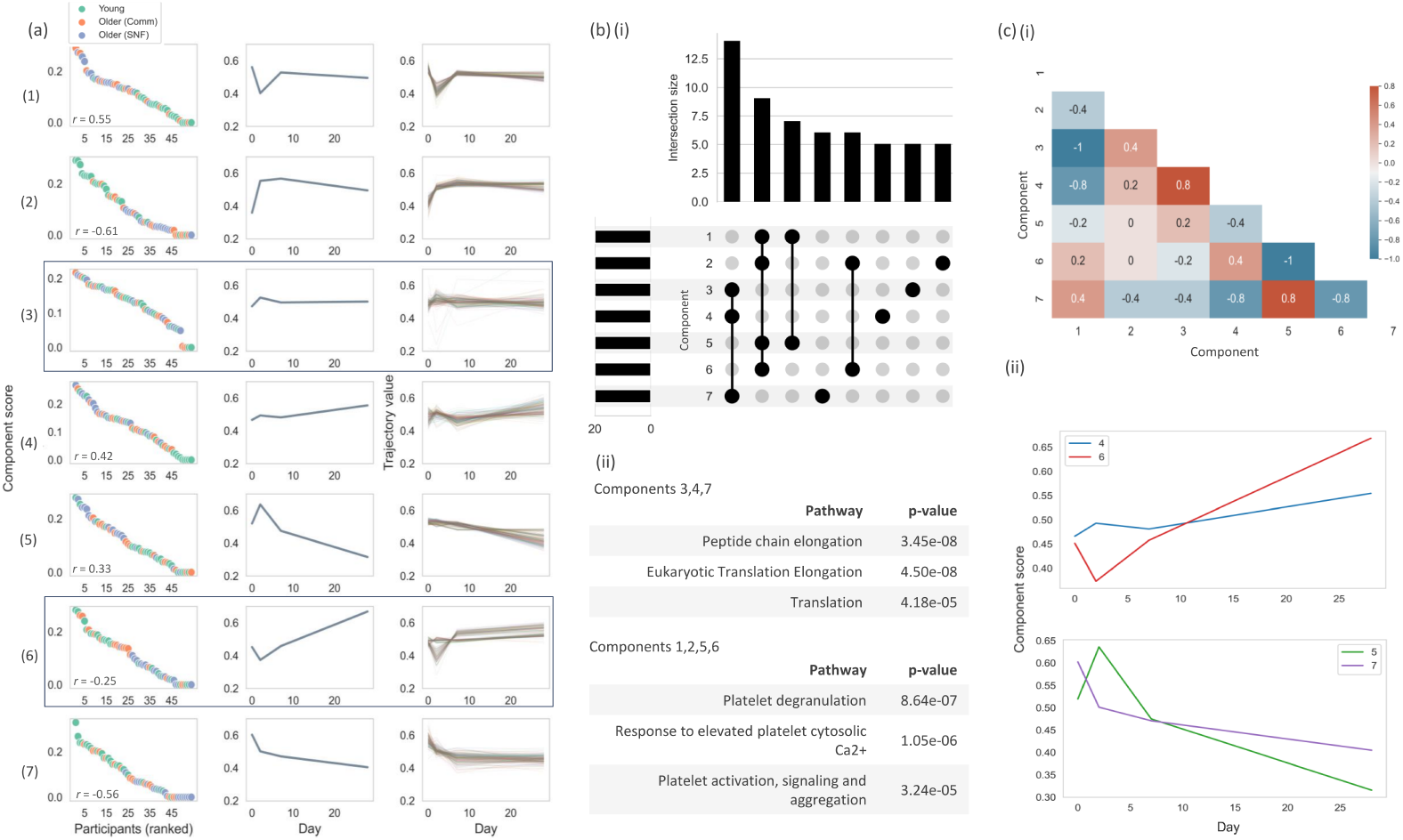
FCT analysis of the selected rank-7 NCPD of platelet transcriptomic data. (a) Participant factor weights, canonical trajectories (CTs), and the top 100 participant–gene trajectories for each component. Reported *r* values indicate significant correlations between participant weights and age; boxes indicate newly identified components. (b) Identification of canonical feature-sets (CFs) using overlap among top-scoring features across components, together with pathway analysis of the largest CFs. (c) Correlations among temporal factor vectors used to identify related CTs, including groups characterized by increasing or decreasing gene expression following vaccination.

These results show that selecting a rank and trial using the QC measures reveals additional, biologically meaningful components that extend prior findings, including new patterns associated with vaccine responsiveness.

We further interpret these results through the FCT framework by examining how the identified components can be expressed in terms of recurring CTs and CFs.

Although several components have distinct temporal patterns, components 4 and 6 can be viewed, at a coarser resolution, as sharing an increasing CT, while components 5 and 7 can be viewed as sharing a decreasing CT (Figure 6c). These shared CTs are paired with different CFs and participant-weight patterns. In component 4, genes associated with translation and peptide elongation increase primarily in older and SNF participants, whereas in component 6, platelet activation genes increase primarily in young and a subset of older participants. Conversely, component 5 captures a decrease in platelet activation genes in older and SNF participants, while component 7 captures a decrease in translation genes primarily in younger participants. This result provides a clear example of how the FCT framework organizes complex biological patterns across participant groups using a smaller number of recurring building blocks derived from the NCPD.

## 3 Discussion

In this work, we developed the Feature Canonical Trajectory framework for modeling longitudinal omics data from the participant perspective. The framework represents each participant’s response as a weighted combination of matrix-valued dictionary elements, or (*Feature Canonical Trajectories*), that are themselves formed from canonical trajectories (CTs) and canonical feature-sets (CFs). We showed that this data model corresponds naturally to the structure of a non-negative CP decomposition (NCPD), and used this connection to derive novel component-level quality-control metrics. These metrics help guide the selection of rank and trial, providing insight beyond global measures that enables the identification of a decomposition with highly interpretable components.

The framework can complement existing tensor decomposition methods developed for longitudinal omics data. For example, the TEMPoral TEnsor Decomposition (TEMPTED) method uses a CP decomposition structure with a functional component to account for a continuous time variable and varying temporal scaling schemes [12]. However, the method does not guarantee that each sample-feature pair will be associated with only one temporal pattern, nor does it provide a mechanism for selecting an optimal rank. The orthogonality and component variance measures introduced in this work could be adapted to the output of the TEMPTED decomposition. The FCT framework could then be used to better interpret the resulting decomposition, with the only difference that the CTs would be represented by continuous functions rather than vectors. A similar approach could be applied to other tensor-based methods for longitudinal omic data, including Tensor Component Analysis (TCAM) [13] and related CP-based approaches that incorporate temporal structure, to improve both decomposition selection and interpretability.

The component variance metric can be understood by examining how well the canonical trajectory (CT) associated with each component represents the trajectories that contribute most strongly to that component. This perspective differs from approaches that operate on reshaped data, such as flattening the tensor into a time × (participant × feature) matrix and clustering time-course trajectories. While such approaches can group trajectories according to similarity, the resulting clusters are defined independently, and therefore do not directly provide a mechanism for relating participants or features across trajectory groups without additional steps. In contrast, the rank-one structure of the NCPD defines components that simultaneously capture coordinated patterns across participants, features, and time. When viewed through the FCT framework, this yields an interpretable representation in which each participant is expressed as a weighted combination of Feature Canonical Trajectories (FCTs), and these weights can be used to group similar participants (and, analogously, features).

This formulation also extends naturally to higher-order tensors arising in more complex study designs. For example, studies such as COMBAT and the IMMuno Phenotyping Assessment in a COVID-19 Cohort (IMPACC) generated multi-tissue data across multiple time points from COVID-19-infected participants, yielding participant × tissue × feature × time tensors for each omic dataset [1, 14]. The NCPD decomposition generalizes to tensors with any number of modes, and the FCT framework can be extended accordingly by introducing additional canonical units associated with each mode.

More broadly, the underlying strategy can be generalized beyond immune profiling studies to other experimental designs. For example, studies measuring feature levels across cells, cell types, or individuals under multiple experimental or physiological conditions naturally yield sample × feature × condition tensors. Such designs arise in multi-tissue, multi-participant expression [6, 15] studies and in single-cell analyses aggregated into sample × cell-type × feature tensors [16, 17]. In this setting, one can view each sample slice of the tensor as a sum of rank-one matrices formed from the outer product of CFs and condition ‘modules’ - sets of conditions under which CFs are up- or down-regulated. While the component variance and orthogonality calculations would need to be adjusted for this particular data model (Supplementary Material), they could be used in an analogous manner to identify a rank and trial of an NCPD that yields interpretable components.

Another established measure of CP decomposition quality is CORCONDIA, introduced by Bro and Kiers [18], which evaluates the structural *appropriateness* of the CP model by assessing how well the factor matrices regress onto a super diagonal core tensor in a corresponding Tucker decomposition. The method requires each factor matrix to be full rank, and its proposed modifications for rank-deficient cases may not be applicable in settings where real-world factors are highly colinear, as is often the case in biological datasets and is accommodated within the FCT framework. Moreover, a decomposition that is well-described by a CP model may still lack alignment with the underlying data model, thereby limiting component-level interpretability. In contrast, the orthogonality and component variance metrics introduced in this work are designed to assess this alignment directly, evaluating whether components are both well-separated and internally consistent with respect to the trajectories they are intended to represent within the FCT framework.

The current limitations of the framework and quality-control metrics are as follows. While the orthogonality and component variance scores provide useful information for selecting a rank and trial, they should be used in conjunction with other criteria when choosing a decomposition for downstream analysis. Global measures such as reconstruction error and similarity provide complementary information regarding model fit and stability that is not captured by component-level measures alone. In addition, as observed in the synthetic data analysis, the FCT framework is not unique, i.e. different combinations of CFs and CTs can yield decompositions with similarly low orthogonality and component variance. In such cases, the overall information recovered across components may be comparable even though the individual components differ.

Constructing stable libraries of CFs and CTs across decompositions would therefore require additional constraints on the decomposition or post-processing approaches designed to identify recurring patterns across trials.

One can consider modifying the NCPD objective function to directly enforce orthogonality conditions. Indeed, in Afshar et al. [19], an orthogonality condition was added to the NCPD objective function to yield a constrained optimization problem that promoted orthogonality between all components of a decomposition. We do not pursue this direction here, as such constraints would be specific to a given data model and may not generalize across experimental settings. By articulating our data model via a tensor dictionary formulation and associated quality control metrics that are generalizable to other experimental settings, we provide an approach that allows users to take advantage of the well-studied NCPD objective function and implementation and to generate decompositions that closely match their specific data models and yield interpretable structure.

## 4 Methods

### 4.1 Formal derivation of the FCT framework

Here, we present the formal derivation of the FCT framework as an NCPD generative model. We assume that there is a set of k CTs,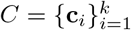 and a set of s CFs, 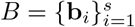. Each CF **b**_*i*_ and CT **c**_*i*_ can combine to form a unique FCT,

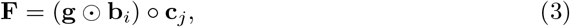

where ⊙ is the Hadamard product and **g** is a vector of scalar weights with the same boolean pattern as **b**i but specific to the FCT.

To index the FCTs for a given decomposition, we observe that the k CTs and s CFs can have a maximum of ks unique combinations, although not all will be realized in a real-world decomposition. For a decomposition of rank *R* ≤ *ks*, we use the indexing operator U : [*R*] → [*k*] ⊗ [*s*], where U(*r*) = (*i, j*) (Figure X). For each r ∈ [*R*], we can denote 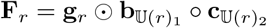 . To simplify the expression, we take 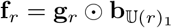 and obtain the following expression for the *r* th FCT,

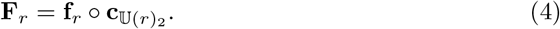

Each participant i in *X* corresponds to a horizontal gene-by-time slice ***X***_*i*_ = *squeeze*(*X*_*i*::_)^*T*^ and can be considered as a weighted sum of FCTs:

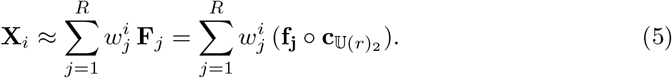

If we now extend this definition to representing all participants, we can write the model as follows:

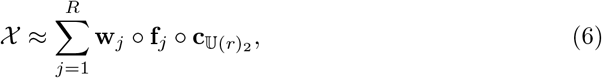

where **w**_*j*_, **f**_*j*_, and 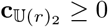. We now observe that Equation 6 represents the data tensor as a sum of rank-one non-negative tensors (components), and thus presents a model of the data in the form of an NCPD (Equation 2) (Figure 2(d)).

### 4.2 Global quality control measures for rank trial and choice

The *normalized Frobenius error* between a data tensor *X* and an NCPD, 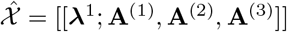 is calculated as

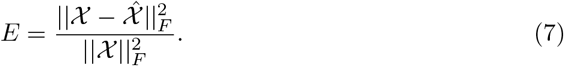

The *similarity score* [5], which is an extension of the triple cosine for third order tensors [20] and Factor Match Score (FMS) [21], quantifies the similarity between two models of the same rank by considering the mean of the weighted product of the cosines of matching pairs of loading vectors, where a match is considered as the maximum score across the set all permutations. For two decompositions of third-order tensors of rank 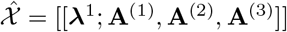 and *ŷ* = [[**λ**^2^ ; **B**^(1)^, **B**^(2)^, **B**^(3)^]], the Similarity,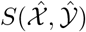, is calculated as

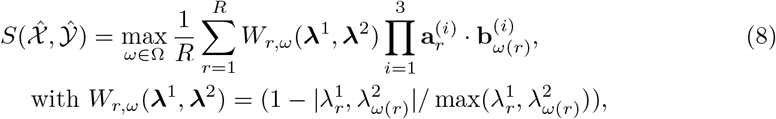

where · is the dot product, 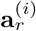 corresponds to the *r* th column of **A**^*(i)*^, and Ω is the set of all permutations of the *R* factors. Note that W penalizes a difference in weights between two components, especially when the weights are of large magnitude. Thus, W penalizes two models that may have otherwise similar components but have been weighted differently.

### 4.3 Component-level quality control measures for rank and trial choice

We introduce two new component-level measures to evaluate trial quality.

#### 4.3.1 Tensor vectorization and index mappings

We define operations to move between tensors and vectors following [22]. To move from a tensor to a vector, we consider a map between a tuple index and a linear index.

##### Definition 6

*For a d-way tensor* 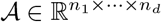, *define*

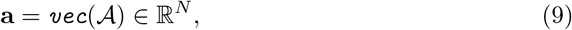

*where* 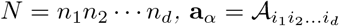, *and α* = L (*i*_1_, *i*_2_, *· · · i*_*d*_). *The operator* L : [*n*_1_] ⊗ [*n*_2_] ⊗ *· · ·* ⊗ [*n*_*d*_] → [*n*_1_*n*_2_ *· · · n*_*d*_] *is given by*

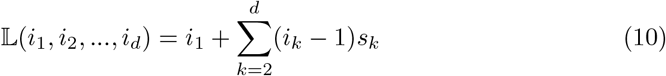

*where* 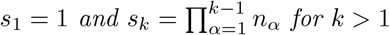.

We also define the operator to move from a vector back to a tensor.

Consider (*i*_1_, *i*_2_, .., *i*_*d*_) = T_*α*_, with T : [*n*_1_*n*_2_ *· · · n*_*d*_] → [*n*_1_] ⊗ [*n*_2_] ⊗ *· · ·* ⊗ [*n*_*d*_] given by

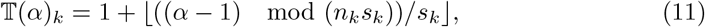

*Where* s1 = 1 and 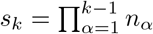 for *k* > 1.

We note that in the case of a tube fiber of *X* ∈ ℝ*n*^*×m×l*^, *X*_*ij*:_ ∈ ℝ ^1*×*1*×l*^, our earlier definition for vec (Equation 1) is consistent with Equation 9, i.e. we take *X*_*ij*:_ = `vec`(*X*_*ij*:_) ∈ ℝ^*l*^ where **x**_*ijk*_ = *X*_*ijk*_ since for **x**_*ij*_*α, α* = *L*(1, 1, *k*) = *k* by Equation 10.

#### 4.3.2 {1, 2}-orthogonality

In {1, 2}-orthogonality, the aim is to assess whether any two components are mutually two-orthogonal in a decomposition. A lower score indicates that a participant and feature will not be assigned to multiple CTs by the NCPD, which would reduce component-level interpretability.

Two rank-one tensors *X* and *Y* are defined as orthogonal if the inner product between any of their constituent vectors is zero (i.e. if any of the constituent vectors are orthogonal to each other) [23]. We extend this definition such that one can specify which subset of constituent factors are required to exhibit orthogonality.

##### Definition 7

*Consider X* = **a**^(1)^ ° **a**^(2)^ ° **a**^(3)^ *and Y* = **b**^(1)^ ° **b**^(2)^ ° **b**^(3)^, *and take* ||a^(i)^|| = ||b^(i)^|| = 1 *for* i ∈ 1, 2, 3, *then X and Y are defined to be S-orthogonal if*

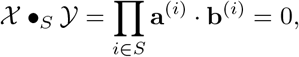

*where* S ⊆ {1, 2, 3}.

If *S* = {1, 2} and *X* •_S_ *Y* = 0, we label *X* and *Y* as {*1,2*}*-orthogonal*. In this case, either the first or second factors are orthogonal in each pair of components. For the decomposition of a participant × gene × time tensor, {1,2}-orthogonality between any two components ensures that for each participant and gene, the associated trajectory is assigned to only one CT. Indeed, it can be shown that if (*X* •_S_ *Y*) = 0, then for any indices (i, j, k) such that *X*_*i,j,k*_≠ 0, *Y*_*i,j,k*_ = 0 (Supplementary Materials).

Following [19], we consider a rank *R* NCPD of a third order tensor 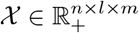 as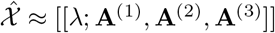, where each of **A**^(*i*)^ are factor matrices of dimension n, l, or m by *R* and are normalized to have unit column vectors. We define the jth component of the decomposition as the rank-one tensor 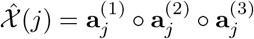, where 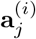 corresponds to the jth column of **A**^(*i*)^.

We now observe that if we take

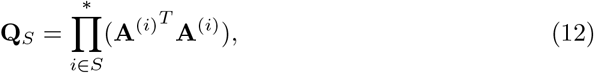

where 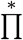 represents a Hadamard product, then **Q***S* is an *R*× *R* matrix where the *i, j*th entry is 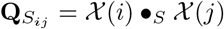. We note that an NCPD will have *S*-orthogonality between all components if **Q**_S_ = **I**_*R*×*R*_ (where **I** is the identity matrix). Therefore, we can assess how far an NCPD is from having components mutually *S*-orthogonal by evaluating the off-diagonal elements of **Q**_*S*_. We consider the following metrics to measure the deviance of the off-diagonal from zero elements:

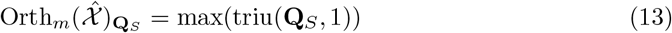

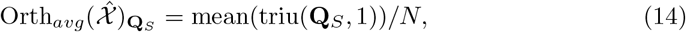

where **Q**_*S*_ is defined in Equation 12, triu(**Q**_*S*_, 1) is a matrix of the upper triangular elements of **Q**_*S*_ without the diagonal, and *N* = (*R*^2^ − *R*)*/*2 is the maximum number of non-zero elements in triu(**Q**_*S*_, 1). We note that in the case of NCPD, **Q**_*s*_ is non-negative, but the definition can be modified to take the absolute value of **Q***s* for an un-constrained CP decomposition.

For the purpose of evaluating the NCPD of a participant *×* gene *×* time data, we take *S* = *{*1, 2*}*. The motivation for this decision is as follows: consider a data tensor *X* of participants *×* genes *×* time, where the gene expression of participants is tracked over discrete time points. The NCPD, 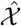, will result in components 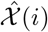 that represent the assignment of certain genes (non-zero values of 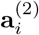, corresponding to a CF) that are expressed in specific individuals indicated by non-zero values in 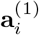 with CT vec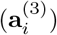. Thus, if a gene *j* for person *k* is assigned a non-zero value to trajectory 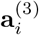, then it should not be assigned to any other trajectory, i.e. the matrix 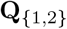 should be as near to **I** as possible, without any assumption on the orthogonality of **A**^(3)^. The orthogonality results in the main text correspond to 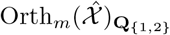 and 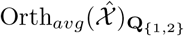, respectively, following these considerations.

#### 4.3.3 Component Variance

The measurement of *component variance* is motivated by the following observation: the participant–gene trajectories that contribute most strongly to a component should be similar to the canonical trajectory (CT) represented by that component. A large distance between these high-weight trajectories and the corresponding CT indicates that the CT does not provide an appropriate representation of the data assigned most strongly to the component. To identify the high-weight trajectories for component *i*, we form the outer product of the corresponding participant and feature factor vectors, vectorize the resulting matrix, and sort its entries in decreasing order. We define the *component variance* 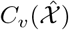 of a rank-*R* decomposition 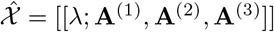 as

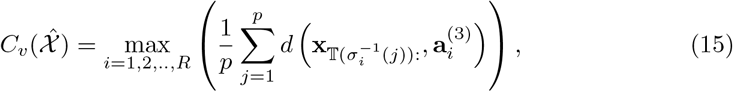

where, for each component *i, σi* is the permutation that arranges the entries of `vec`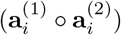 in decreasing order, where 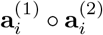 is the outer product of the participant factor vector and the feature factor vector representing the CF for component *i*. The same permutation is applied to the participant–gene trajectories in the data tensor, so that 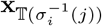 denotes the observed trajectory at the participant–gene index corresponding to the *j*th-largest entry of `vec`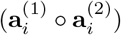. Here, *d* is a user-specified distance function used to compare an observed participant–gene trajectory with the corresponding CT. The mapping *T* : *[nl] → [n] ⊗ [l]* maps each vectorized index back to its corresponding participant–gene index, as defined in Equation 11. The threshold *p* specifies the number of highest-weight participant–gene trajectories included in the analysis.

#### 4.3.4 Using global and component-level QC measures to select a rank and trial

Global and component-level QC measures provide complementary information about decomposition quality. A decomposition with low component variance but high *{*1, 2*}*-orthogonality may contain components whose highest-weight trajectories are well represented by their associated CTs, but whose participant–feature assignments overlap across components. Conversely, a decomposition with high component variance but low *{*1, 2*}*-orthogonality may assign each participant–feature trajectory primarily to a single component, while at least one component CT does not adequately represent the trajectories assigned to it. A decomposition with relatively high reconstruction error but low component variance and *{*1, 2*}*-orthogonality may be missing one or more components, even though the components it recovers are interpretable. Finally, a decomposition with low reconstruction error but high component variance or *{*1, 2*}*-orthogonality may fit the data globally while containing components with poor internal consistency or separation. Therefore, multiple trials should be evaluated at each rank, and rank and trial selection should take into account both global and component-level QC measures.

### 4.4 Tensor decomposition

The NCPD was performed using *all-at-once* optimization (CP-OPT) with a non-negative lower bound in the Tensor Toolbox package [24]. CP-OPT is a gradient-based optimization method that has been shown to be more accurate than CP-ALS (alternating least squares) and more efficient than CP-NLS (nonlinear alternating least squares) [25]. We use CP-OPT with the objective function 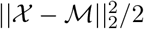, where *X* is the input data, *M* is the reconstructed model, and || *·* ||_2_ is the Frobenius norm. The decomposition was initialized using random conditions, and performed with a range of ranks and multiple trials per rank. The quality of the decompositions was evaluated using the normalized Frobenius error, similarity score, *{*1, 2*}*-orthogonality score, and component variance. The decomposition models chosen for evaluation were subsequently analyzed using the FCT framework.

### 4.5 Datasets

#### 4.5.1 Synthetic data

In the synthetic dataset, we consider that gene expression post-vaccination can correspond to one of three CTs. The genes themselves correspond to three CFs: immune, metabolic, and house-keeping. Finally, transcription data is collected from three groups of participants: healthy young (Y), healthy older (O), and immunocompromised of any age (IC) (Figure 2(a)).

From the three CFs, and three CTs, we construct six FCTs (Fig 2(b)). For example, **F**_1_ is the outer product of the immune gene CF and the rise-and-fall CT, which represents a healthy immune response. Each group of participants is composed of a unique combination of FCTs (Fig. 2(c)). For instance, the Older group shows a decrease in immune CF activity over time (**F**_3_), a rise-and-fall of metabolic gene activity (**F**_4_), and no change in house-keeping gene activity (**F**_6_). To introduce heterogeneity within each canonical feature-set, gene-specific weights were independently sampled for each gene–FCT pair. The resulting weighted CFs were then combined with the CTs via outer products to form the FCTs. We construct the components as the outer product between the associated participant weight vector, CF, and CT for FCTs 1-6. The final synthetic tensor is the sum of the components (Fig. 2(d)).

#### 4.5.2 Real-world data

Transcriptomic data derived from platelets of individuals vaccinated with the influenza vaccine as previously described in [11] was used. Briefly, young adults (age 21–35 years), community-dwelling older adults (age ≥ 65 years) (Older (Comm)), and older adults (age ≥ 65 years) (Older (SNF)) who were residents of a skilled nursing facility (SNF) in greater New Haven, Connecticut were enrolled. All participants received the seasonal high-dose influenza vaccine. Platelet-rich plasma (PRP) was isolaged from blood samples of participants obtained prior to vaccination (Day 0) and at Days 2, 7, and 28 post-vaccine for isolation of RNA and elucidation of the platelet transcriptome via RNA-seq. The data was pre-processed to only include protein-coding genes and exclude genes on the X and Y chromosome. Following, genes were filtered to remove genes with counts ≤ 100 in ≥ 30 samples (which corresponds to approximately 12% of total samples), and the bottom 10% expressing genes were then filtered from the remainder. After filtering, gene counts were normalized using DESeq2 median of ratios method, and log2(*x* + 1) transformed [26]. The 500 most variable genes in the count space were used for the decomposition in a tensor framework of participant *×* gene *×* day. Only participants with data for all days in the study were used. The final dimensions of the tensor were 54 *×* 500 *×* 4. Vaccine responsiveness was assessed using hemagglutination inhibition (HAI) antibody titers and reflects the magnitude of the antibody response to influenza vaccination; participants were classified as high or low responders based on their post-vaccination serological response as described in [11].

## 5 Code availability

Code used to generate the synthetic decompositions and compute the quality-control metrics presented in this work is publicly available at https://github.com/akonstodata/FCT. The repository includes implementations of the component variance and *{*1, 2*}*-orthogonality metrics, together with example workflows for both Tensor Toolbox and TensorLy decompositions.

## 6 Acknowledgments

The authors would like to acknowledge Shaoib Bin Masud and Eric Phipps for helpful discussions regarding the manuscript. S.H.K., M.K., A.K., and J.X. would like to acknowledge support of the National Institutes of Health grant R21AI176204.

