## Supplementary Materials for "Assessing tensor decomposition quality of immune profiling data from a dictionary learning perspective"

#### S1 On the non-uniqueness of the FCT framework

The non-uniqueness of the FCT representation can be illustrated using the synthetic immune-response dataset. This example also shows that the underlying Canonical Feature-sets (CFs) can still be recovered through post-processing when desired. In the synthetic immune response dataset (Fig. 2, main text), there are three FCTs ($\mathbf{F}_2$, $\mathbf{F}_5$, and $\mathbf{F}_6$) that share the same constant Canonical Trajectory (CT) ($\mathbf{c}_2$) taken as an outer product with different CFs ($\mathbf{g}_1$, the immune gene CF; $\mathbf{g}_2$, the metabolic gene CF, and $\mathbf{g}_3$, the house-keeping gene CF, respectively). Only the immunocompromised group of participants has non-zero weights for $\mathbf{F}_2$, since only in immunocompromised participants do the immune genes not change upon vaccination; healthy young and immunocompromised participants have non-zero weights for $\mathbf{F}_5$, since metabolic genes only change in the older group upon vaccination; and all participants have non-zero weights for $\mathbf{F}_6$, as house-keeping genes do not change over time for all participants.

The three ground truth components, as well as three components from an NCPD of rank $R = 6$ with lowest error and component variance out of 100 trials, corresponding to the constant CT $\mathbf{c}_2$, are visualized in the main text Fig. 4 (a-b) and Supplementary Fig. 1. We observe that the NCPD identifies the same FCT as the ground truth characterized by taking the metabolic gene CF $\mathbf{g}_2$ as an outer product with the constant CT ($\mathbf{F}_5 = \mathbf{g}_2 \circ \mathbf{c}_2$) (Supplementary Fig. 1), with non-zero values for young and immuno-compromised participants in both. If we take $\widehat{\mathcal{Y}} = [[\lambda, \mathbf{A}, \mathbf{B}, \mathbf{C}]]$ as the ground truth decomposition, and $\mathcal{X} = [[\gamma, \mathbf{A}*, \mathbf{B}*, \mathbf{C}*]]$ as the NCPD of rank $R = 6$ as noted above, then the above statement can be rewritten as $\mathbf{a}_4 \circ \mathbf{b}_4 \circ \mathbf{c}_4 \approx \mathbf{a}*_4 \circ \mathbf{b}_4 \circ \mathbf{c}_4$, where 4 corresponds to the 4th component of each respective decomposition.

Nevertheless, we observe that the original $\mathbf{F}_2$ and $\mathbf{F}_6$ (corresponding to ground-truth components 5 and 6) are re-organized in the NCPD (Fig. 4 (a-b)). While the decomposition still contains an FCT with the house-keeping gene CF taken as an outer product with the constant CT (component 6 of the decomposition), note that unlike in the ground-truth component, only healthy young and older participants are strongly associated with this FCT, whereas in the original, all participants are.

This apparent difference is resolved by examining component 5 of the fitted NCPD. Here, we observe both the house-keeping and immune gene CFs grouped together into one FCT, and are associated with just the immuno-compromised participants. From a linear algebraic perspective, the sum of these two components yields the same result (Supplementary Fig. 2). For simplicity, let $\mathbf{w}_{(\mathrm{Y})}, \mathbf{w}_{(\mathrm{O})}, \mathbf{w}_{(\mathrm{I})} \in \mathbb{R}+^{15}$ denote participant-weight vectors with nonzero entries only for the healthy young, healthy older, or immunocompromised participants, respectively. We write $\mathbf{w}_{(\mathrm{YOI})} = \mathbf{w}_{(\mathrm{Y})} + \mathbf{w}_{(\mathrm{O})} + \mathbf{w}_{(\mathrm{I})}$ for the participant-weight vector with nonzero entries across all three groups. For this illustrative equivalence, we assume that the nonzero weights on the participants are approximately the same in the relevant components. Under this simplifying assumption, the equivalence can be written as

$$\mathbf{w}_{(\mathrm{I})} \circ \mathbf{g}_1 \circ \mathbf{c}_2 + \mathbf{w}_{(\mathrm{YOI})} \circ \mathbf{g}_3 \circ \mathbf{c}_2 \tag{S1}$$

$$= \mathbf{w}_{(\mathrm{I})} \circ \mathbf{g}_1 \circ \mathbf{c}_2 + \mathbf{w}_{(\mathrm{I})} \circ \mathbf{g}_3 \circ \mathbf{c}_2 + \mathbf{w}_{(\mathrm{YO})} \circ \mathbf{g}_3 \circ \mathbf{c}_2$$

$$= \mathbf{w}_{(\mathrm{I})} \circ (\mathbf{g}_1 + \mathbf{g}_3) \circ \mathbf{c}_2 + \mathbf{w}_{(\mathrm{YO})} \circ \mathbf{g}_3 \circ \mathbf{c}_2. \tag{S2}$$

Note that Equation S1 corresponds to the ground truth components 5 and 6, and Equation S2 corresponds to components 5 and 6 of the fitted NCPD (Fig. 4 (a,b), main text; Supplementary Figure 2 (b)). Observe that in Equation S1, $\mathbf{g}_1 \cdot \mathbf{g}_3 = 0$ and in Equation S2, $\mathbf{w}_{(\mathrm{I})} \cdot \mathbf{w}_{(\mathrm{YO})} = 0$, therefore the orthogonality of the two components in each decomposition is preserved (see Section S2 for further details).

Since the NCPD optimization algorithm does not constrain the feature factor to be orthogonal, it can identify algebraically equivalent combinations of FCTs as discussed above. Although orthogonality constraints could be imposed directly on the feature factors, unique CFs can also be recovered through a post-processing intersection analysis while retaining the standard NCPD objective. Note that the biological patterns revealed by both the ground-truth and NCPD in this example are equivalent, therefore the non-uniqueness of the framework does not preclude the same downstream analysis and interpretation of the underlying system.

#### S2 $\{1,2\}$-orthogonality

We prove that if two non-negative NCPD components are $\{1,2\}$-orthogonal, then they cannot both have a nonzero value at the same tensor entry. Consequently, a participant–gene trajectory can be assigned to at most one of the two components. We first review two definitions for a rank-one tensor, and a tensor inner product for two rank-one tensors. Here, we focus on third-order tensors, but the definitions and results can be generalized to a higher order in a straightforward way. We begin by defining a third-order rank-one tensor as follows [4],

**Definition S1.** *A third-order tensor* $\mathcal{X} \in \mathbb{R}^{n\times l\times m}$ *is rank one if it can be written as an outer product of three vectors,* $\mathcal{X} = \mathbf{a}^{(1)} \circ \mathbf{a}^{(2)} \circ \mathbf{a}^{(3)}$*, where* $\mathbf{a}^{(1)} \in \mathbb{R}^n, \mathbf{a}^{(2)} \in \mathbb{R}^l, \mathbf{a}^{(3)} \in \mathbb{R}^m$*.*

We define two rank-one tensors to be S-orthogonal when at least one pair of corresponding factor vectors indexed by S is orthogonal.

**Definition S2.** *Let* $\mathcal{X} = \mathbf{a}^{(1)} \circ \mathbf{a}^{(2)} \circ \mathbf{a}^{(3)}$ *and* $\mathcal{Y} = \mathbf{b}^{(1)} \circ \mathbf{b}^{(2)} \circ \mathbf{b}^{(3)}$*. For a nonempty set* $S \subseteq \{1,2,3\}$*,* $\mathcal{X}$ *and* $\mathcal{Y}$ *are S-orthogonal if*

$$\mathcal{X} \bullet_S \mathcal{Y} = \prod_{r\in S} \left(\mathbf{a}^{(r)} \cdot \mathbf{b}^{(r)}\right) = 0.$$

When $S = \{1,2,3\}$, this definition corresponds to the definition of orthogonality of two rank-one tensors in [3].

We show that if two non-negative rank-one third-order tensors are $S-$orthogonal, then they are also orthogonal when reshaped as vectors.

**Proposition S1.** *For two rank-one third-order tensors* $\mathcal{X} = \mathbf{a}^{(1)} \circ \mathbf{a}^{(2)} \circ \mathbf{a}^{(3)}$ *and* $\mathcal{Y} = \mathbf{b}^{(1)} \circ \mathbf{b}^{(2)} \circ \mathbf{b}^{(3)}$*, where* $\mathbf{a}^{(1)}, \mathbf{b}^{(1)} \in \mathbb{R}^n_+, \mathbf{a}^{(2)}, \mathbf{b}^{(2)} \in \mathbb{R}^l_+$*, and* $\mathbf{a}^{(3)}, \mathbf{b}^{(3)} \in \mathbb{R}^m_+$*, if* $\mathcal{X} \bullet_S \mathcal{Y} = 0$ *for a given nonempty* $S \subseteq \{1,2,3\}$*, then* $\texttt{vec}(\mathcal{X}) \cdot \texttt{vec}(\mathcal{Y}) = 0$*, where* $\texttt{vec}(\mathcal{X}) \in \mathbb{R}^{nml}$ *is the vectorized form of* $\mathcal{X}$*.*

*Proof.* Since $\mathcal{X}_{ijk} = a_i^{(1)} a_j^{(2)} a_k^{(3)}$ and $\mathcal{Y}_{ijk} = b_i^{(1)} b_j^{(2)} b_k^{(3)}$, we have

$$\begin{aligned}
\texttt{vec}(\mathcal{X}) \cdot \texttt{vec}(\mathcal{Y}) &= \sum_{i=1}^n \sum_{j=1}^l \sum_{k=1}^m \mathcal{X}_{ijk}\mathcal{Y}_{ijk} \\
&= \sum_{i=1}^n \sum_{j=1}^l \sum_{k=1}^m \left(a_i^{(1)} a_j^{(2)} a_k^{(3)}\right) \left(b_i^{(1)} b_j^{(2)} b_k^{(3)}\right) \\
&= \sum_{i=1}^n \sum_{j=1}^l \sum_{k=1}^m (a_i^{(1)} b_i^{(1)})(a_j^{(2)} b_j^{(2)})(a_k^{(3)} b_k^{(3)}) \\
&= \left(\sum_{i=1}^n a_i^{(1)} b_i^{(1)}\right) \left(\sum_{j=1}^l a_j^{(2)} b_j^{(2)}\right) \left(\sum_{k=1}^m a_k^{(3)} b_k^{(3)}\right) \\
&= \prod_{r=1}^3 \left(\mathbf{a}^{(r)} \cdot \mathbf{b}^{(r)}\right).
\end{aligned}$$

Because $\mathcal{X} \bullet_S \mathcal{Y} = 0$, there exists some $s \in S$ such that $\mathbf{a}^{(s)} \cdot \mathbf{b}^{(s)} = 0$. Therefore, one factor in the product above is zero, and hence $\texttt{vec}(\mathcal{X}) \cdot \texttt{vec}(\mathcal{Y}) = 0$.

□

Proposition S1 tells us that in an NCPD, if two components $\mathcal{X}$ and $\mathcal{Y}$ are $S$-orthogonal for some nonempty $S \subseteq \{1, 2, 3\}$. Since NCPD components are non-negative, this implies that for all indices $(i, j, k)$, if $\mathcal{X}_{i,j,k} \neq 0$, then $\mathcal{Y}_{i,j,k} = 0$. We also show that the reverse is true, i.e. if the vectorized form of components $\mathcal{X}$ and $\mathcal{Y}$ of an NCPD is orthogonal, then $\mathcal{X}$ and $\mathcal{Y}$ are $S$-orthogonal for $S = \{1, 2, 3\}$, indicating that there exists at least one pair of factor vectors that are orthogonal between the two components.

**Corollary S1.** *For two rank-one third-order tensors $\mathcal{X} = \mathbf{a}^{(1)} \circ \mathbf{a}^{(2)} \circ \mathbf{a}^{(3)}$ and $\mathcal{Y} = \mathbf{b}^{(1)} \circ \mathbf{b}^{(2)} \circ \mathbf{b}^{(3)}$, where $\mathbf{a}^{(1)}, \mathbf{b}^{(1)} \in \mathbb{R}^n_+$, $\mathbf{a}^{(2)}, \mathbf{b}^{(2)} \in \mathbb{R}^l_+$, and $\mathbf{a}^{(3)}, \mathbf{b}^{(3)} \in \mathbb{R}^m_+$, if $\texttt{vec}(\mathcal{X}) \cdot \texttt{vec}(\mathcal{Y}) = 0$, where $\texttt{vec}(\mathcal{X}) \in \mathbb{R}^{nml}$ is the vectorized form of $\mathcal{X}$, then $\mathcal{X} \bullet_S \mathcal{Y} = 0$ for $S = \{1, 2, 3\}$.*

*Proof.* From the calculation in the proof of Proposition S1,

$$\texttt{vec}(\mathcal{X}) \cdot \texttt{vec}(\mathcal{Y}) = \prod_{r=1}^{3} \left( \mathbf{a}^{(r)} \cdot \mathbf{b}^{(r)} \right).$$

Since $\texttt{vec}(\mathcal{X}) \cdot \texttt{vec}(\mathcal{Y}) = 0$, at least one factor in this product must be zero. Therefore,

$$\mathcal{X} \bullet_{\{1,2,3\}} \mathcal{Y} = \prod_{r=1}^{3} \left( \mathbf{a}^{(r)} \cdot \mathbf{b}^{(r)} \right) = 0.$$

□

For a participant × gene × time tensor, Proposition S1 implies that if two components are $\{1, 2\}$-orthogonal, then no participant–gene pair can have nonzero weight in both components. Thus, the trajectory associated with a given participant–gene pair can be assigned to at most one CT across the two components.

### S3 Extension of the FCT framework and metrics for categorical data

While the FCT framework and associated metrics were designed for immune profiling data organized as participants × features × time, it can be readily extended for data structured as individuals × features × experiment. The experiment mode may represent different cell or tissue types measured in the same individuals [1, 5, 6], or different omics assays [2]. For this data model, individuals can be represented as a weighted combination of the outer products of Canonical Feature Sets (CFs) and Canonical Experimental Modules (CEMs). The CEMs replace the Canonical Trajectories (CTs) in the third mode and are defined analogously to the CFs.

The component variance and orthogonality scores are modified accordingly. For component variance, we evaluate the alignment between the highest-weight individual–feature–experiment entries of each component and the corresponding entries of the observed tensor. For data $\mathcal{X} \in \mathbb{R}^{n \times l \times m}_+$ with structure individuals × features × experiments, let $\widehat{\mathcal{X}} = [[\boldsymbol{\lambda}; \mathbf{A}^{(1)}, \mathbf{A}^{(2)}, \mathbf{A}^{(3)}]]$ be a rank-$R$ decomposition. We define the component variance $C_v(\widehat{\mathcal{X}})$ as

$$C_v(\widehat{\mathcal{X}}) = \max_{i=1,\ldots,R} d\left(\mathcal{X}_{\mathbb{T}(\sigma_i^{-1}(1:p))}, \widehat{\mathcal{X}}(i)_{\mathbb{T}(\sigma_i^{-1}(1:p))}\right). \quad \text{(S3)}$$

For each component $i$, $\sigma_i$ is the permutation that arranges the entries of $\texttt{vec}(\widehat{\mathcal{X}}(i))$ in decreasing order. The same permutation is applied to $\texttt{vec}(\mathcal{X})$, so that $\mathcal{X}_{\mathbb{T}(\sigma_i^{-1}(1:p))}$ and $\widehat{\mathcal{X}}(i)_{\mathbb{T}(\sigma_i^{-1}(1:p))}$ denote the vectors of observed and component values, respectively, at the tensor indices corresponding to the $p$ largest entries of component $i$. Here, $d$ denotes a user-specified distance between these two vectors, and $\mathbb{T} : [nlm] \to [n] \otimes [l] \otimes [m]$ maps each vectorized index back to its corresponding individual–feature–experiment tensor index, as defined in Equation 11 of the main text.

The orthogonality score can be modified by taking $S = \{1, 2, 3\}$ in Equations 12–14 of the main text. In this setting, orthogonality in any mode between two components does not compromise interpretability because all three modes are categorical. A rank-$R$ decomposition $\widehat{\mathcal{X}}$ is therefore interpretable under this criterion if each individual–feature–experiment entry is assigned to at most one component. That is, if $\widehat{\mathcal{X}}(r)_{ijk} > 0$, then $\widehat{\mathcal{X}}(s)_{ijk} = 0$ for every $s \in \{1, \ldots, R\}$ with $s \neq r$. By Corollary S1, this condition is equivalent to requiring $\widehat{\mathcal{X}}(r) \bullet_S \widehat{\mathcal{X}}(s) = 0$ for every pair of distinct components $r, s \in \{1, \ldots, R\}$.
