## Supplementary Figures for "Assessing tensor decomposition quality of immune profiling data from a dictionary learning perspective"

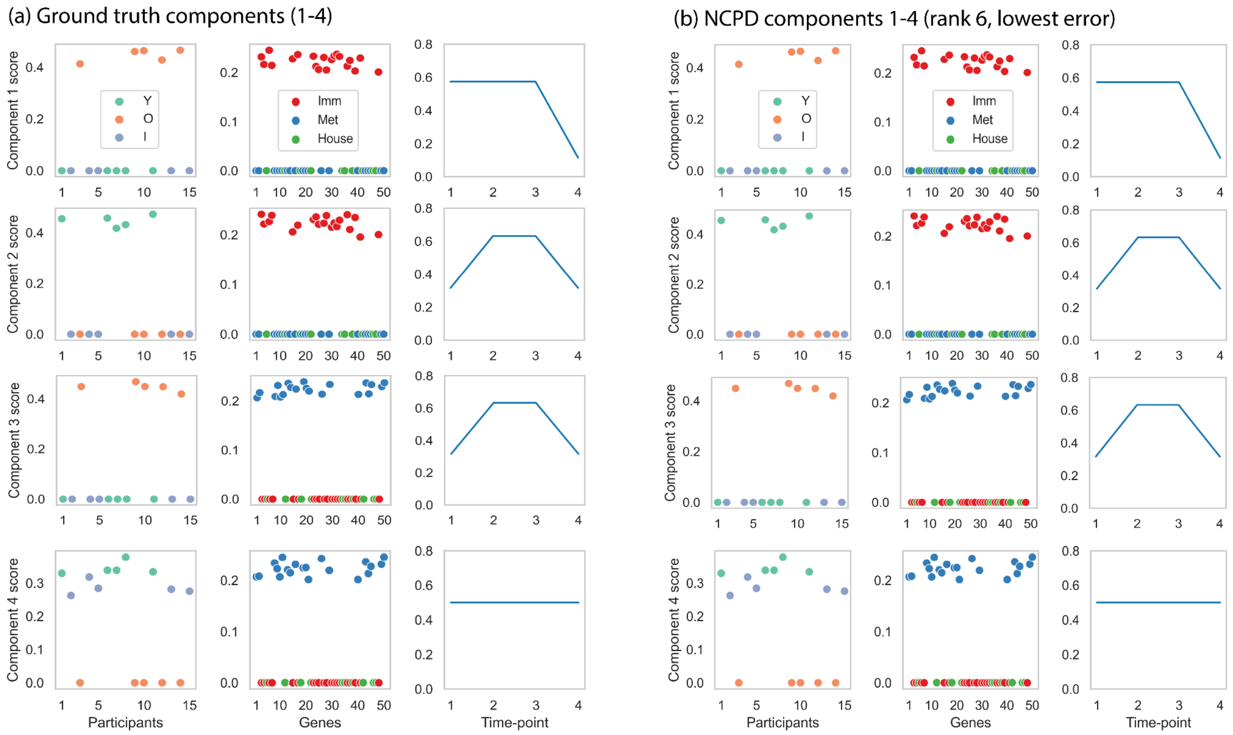


**Supplementary Figure 1** (a) Ground truth components 1-4 and (b) components 1-4 from the rank *R*=6 NCPD trial with the lowest error and component variance for the synthetic dataset.

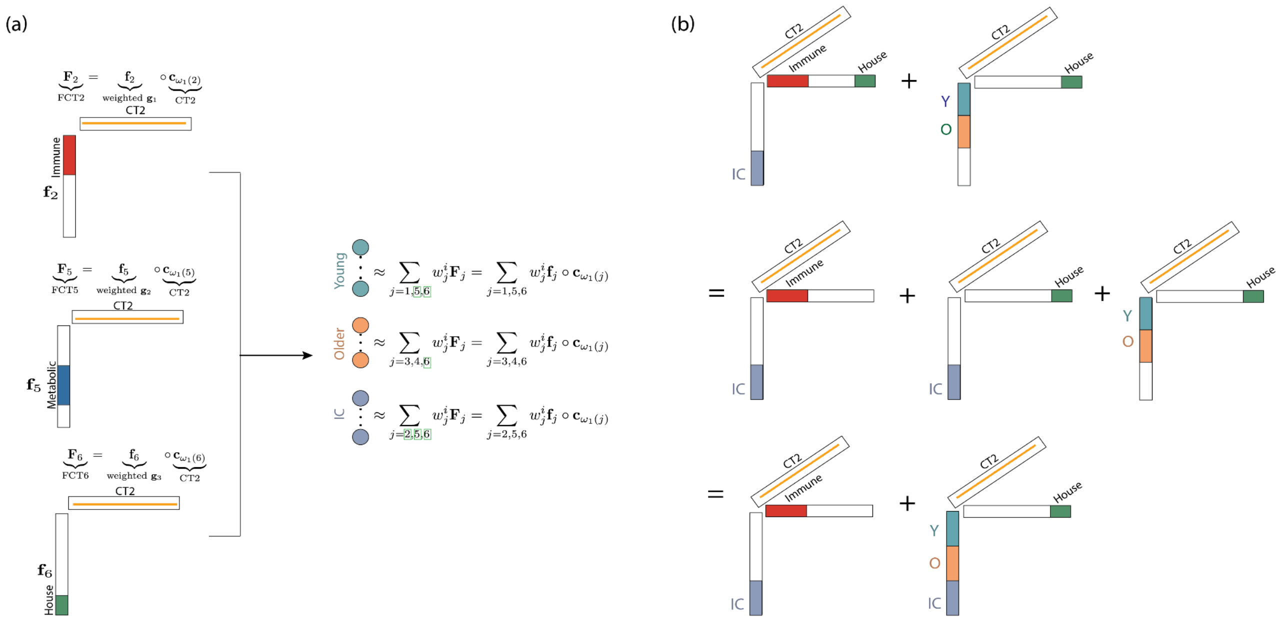


**Supplementary Figure 2** Equivalent representations of the FCT framework for the synthetic dataset. (a) The three ground-truth FCTs associated with the constant Canonical Trajectory CT2 and their contributions to the dictionary representation of healthy young, healthy older, and immunocompromised participants. (b) Algebraic equivalence between the relevant components of the fitted NCPD with the lowest error and component variance (top row) and the corresponding ground-truth components (bottom row).

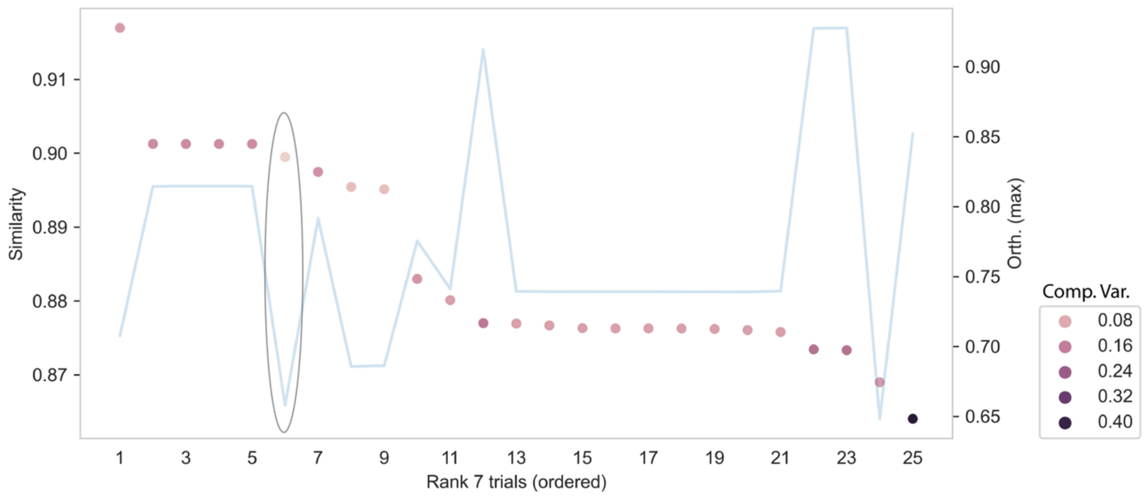


**Supplementary Figure 3** Similarity of rank $R$=7 trials to the rank $R$=6 trial with the lowest component variance of the NCPD of the platelet transcriptomic data. Similarity is shown by the scatter points on the left axis, with point color indicating component variance, and the maximum orthogonality score is shown by the blue line on the right axis. The circled trial ranks sixth in similarity, has the lowest component variance across all trials (0.014), and has the lowest maximum orthogonality score (0.66) among the most similar trials.

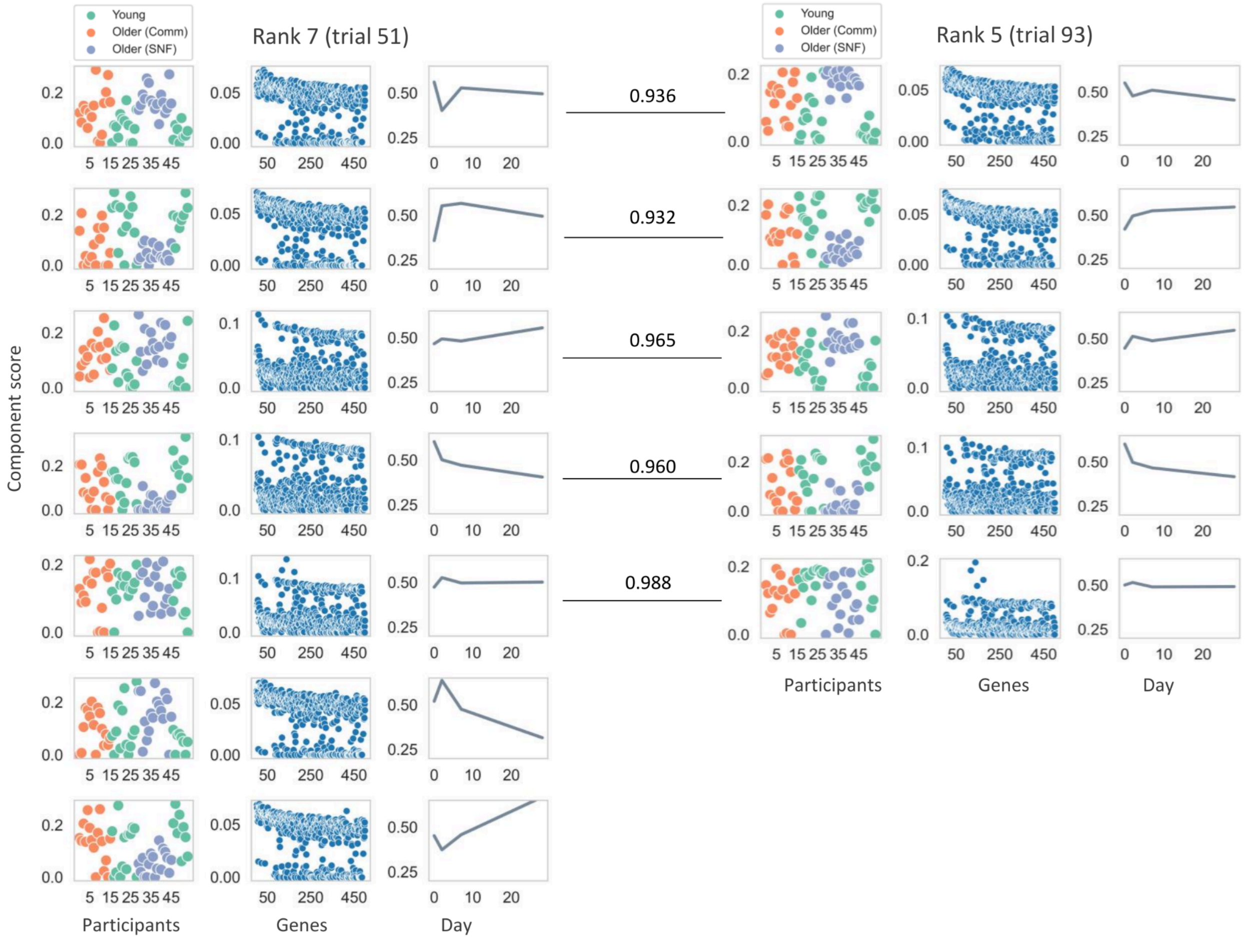


**Supplementary Figure 4** Comparison of the trials with the lowest component variance at ranks *R*=7 and *R*=5 of an NCPD of the platelet transcriptomic data. Components of the rank *R*=7 decomposition (trial 51) were reordered to align with the components of rank *R*=5 (trial 93) with which they have the highest component similarity. Component similarity is defined as the product of the cosine similarities between the corresponding factor vectors of a pair of components.

| Comp. | Correlation (age) | p value<br>MW sex | <br>KW group | <br>MW response |
|---|---|---|---|---|
| 1 | **0.55 (1.49e-05)** | 4.84E-01 | **9.61E-06** | **2.84E-02** |
| 2 | **-0.61 (9.23e-07)** | 5.81E-01 | **1.21E-04** | 4.73E-01 |
| 3 | 0.03 (8.38e-01) | 8.75E-01 | 9.91E-01 | 7.34E-01 |
| 4 | **0.42 (1.62e-03)** | 5.01E-01 | **1.67E-03** | 1.80E-01 |
| 5 | **0.33 (1.62e-02)** | 2.62E-01 | **2.19E-02** | 5.45E-01 |
| 6 | **-0.25 (7.16e-02)** | 1.78E-01 | **8.99E-03** | **3.13E-02** |
| 7 | **-0.56 (1.2e-05)** | 7.19E-01 | **3.16E-05** | **9.18E-02** |

**Supplementary Table 1** Association of component scores with age (Pearson correlation) and group (Kruskal‑Wallis test), biological sex, and vaccine response (Mann‑Whitney U test). Bolded values indicate significance of association ($p < 0.10$).
